# Chronic cardiotoxicity assessment by cell optoporation on microelectrode arrays

**DOI:** 10.1101/2022.06.20.496820

**Authors:** Giuseppina Iachetta, Giovanni Melle, Nicolò Colistra, Francesco Tantussi, Francesco De Angelis, Michele Dipalo

**Affiliations:** Istituto Italiano di Tecnologia, Via Morego 30, 16163, Genova, Italy; FORESEE Biosystems srl, Genova, Italy

**Keywords:** Chronic cardiotoxicity, optoporation, intracellular recording, cardiomyocytes, in-vitro toxicology, action potentials

## Abstract

The reliable identification of chronic cardiotoxic effects in *in vitro* screenings is fundamental for filtering out toxic molecular entities before *in vivo* animal experimentation and clinical trials. Present techniques such as patch-clamp, voltage indicators, and standard microelectrode arrays do not offer at the same time high sensitivity for measuring transmembrane ion currents and low-invasiveness for monitoring cells over long time. Here, we show that optoporation applied to microelectrode arrays enables measuring action potentials from human-derived cardiac syncytia for more than 1 continuous month and provides reliable data on chronic cardiotoxic effects caused by known compounds such as pentamidine. The technique has high potential for detecting chronic cardiotoxicity in the early phases of drug development.

## Introduction

The process of drug discovery is extremely long and expensive, with cost estimates ranging from $314 million to $2.8 billion and a period of 10 to 15 years to commercialize a new drug (Morgan et al. 2011; DiMasi et al. 2016). Considering the development cost and time, the reduction of high attrition rates during this process is still a key challenge for the pharmaceutical companies (Hay et al. 2014). Although reasons for drug attrition and market withdrawals are several, the major cause of drug failure can be associated with their cardiac adverse effects (Ferdinandy et al. 2019).

The current cardiac safety pharmacology guidelines (ICH S7B and E14) mandate *in vitro* measurements of the human-ether-à-go-go Related Gene (hERG) potassium channel activity followed by *in vivo* QT prolongation measurements as traditional standards for preclinical screening assay to evaluate proarrhythmic risk of all candidates drug (FDA 2005a, b). In addition, the Comprehensive *in vitro* Proarrhythmia Assay (CiPA) initiative, established in 2013 to develop a new paradigm for assessing proarrhythmic risk, validated the use of microelectrode array (MEA) and cardiomyocytes derived from human induced pluripotent stem cells (hiPSC-CMs) to assess drug induced cardiotoxicity during the *in vitro* preclinical phase (Colatsky et al. 2016; Kitaguchi et al. 2016; Takasuna et al. 2017; Blinova et al. 2018). In parallel, research groups are evaluating the implementation of 3D nanostructures on MEAs and the development of novel optical approaches to enable recordings of cardiac action potentials (APs) from *in vitro* two-dimensional syncytia (Dipalo et al. 2017; Desbiolles et al. 2020; Hu et al. 2020; Barbaglia et al. 2021; Zhou et al. 2021; Jahed et al. 2022). However, all current methods for *in vitro* cardiotoxicity screenings focus on acute effects and short time-points after drug administration (from minutes to a few hours), missing their possible transient and reversible effects as well as cell responses after long-term administration. Notably, several compounds such as hERG channel trafficking inhibitors (e.g. pentamidine) (Dennis et al. 2007; Obergrussberger et al. 2016; Asahi et al. 2019) and chemotherapeutic molecules (e.g. doxorubicin) (Kumar et al. 2012; Chaudhari et al. 2017; Mladěnka et al. 2018; Narezkina and Nasim 2019; Bozza et al. 2021) may not show their cardiotoxic effects after acute dose administration but may become toxic after repeated and prolonged exposures (several hours to days) (Cai et al. 2019). Thus, adverse cardiac effects occurring after long-term drug exposure may be undetected during the preclinical screening and may allow for the further development of potentially toxic molecules. Although hiPSC-CMs can be maintained in culture for long periods of time with stable baseline functions (Kopljar et al. 2017; Dias et al. 2018), the current methods such as patch-clamp, voltage indicators, and MEA technology do not offer at the same time high sensitivity for measuring transmembrane ion currents and low-invasiveness for monitoring cells over long time. Therefore, despite the recent advances in chronic cardiotoxicity assays using hiPSC-CMs (Narkar et al. 2022), long-term (chronic) cardiac adverse effects remain a major barrier of drug development. The *in vitro* detection of chronic cardiotoxicity represents a key turning point to avoid the progression of a drug candidate with “delayed” cardiac adverse effect. Thus, to render the drug development process more efficient, there is an immediate need to develop standardized assays to detect chronic cardiotoxicity during *in vitro* screenings.

We recently proposed and validated a novel approach for high-quality recordings of intracellular action potentials (APs) simultaneously from large cardiomyocyte networks combining commercial MEA technology with cell optoporation (Dipalo et al. 2018; Melle et al. 2020; Iachetta et al. 2021). Our approach improves the reliability of acute cardiotoxicity drug screenings reducing the cell cultures required to reach exploitable results in cardiac safety assessment.

By exploiting the non-invasiveness of optoporation (Messina et al. 2015) and a new experimental procedure based on backside laser excitation of transparent electrodes, in this work we demonstrate that it enables extremely long-term AP recordings of the same cells on commercial MEAs, paving the way to the reliable chronic cardiotoxicity assessment on hiPSC-CMs. We used optoporation on multiwell 60-electrode commercial MEAs with titanium nitride (TiN) transparent and thin electrodes to measure the APs from the same hiPSC-CMs for up to 35 *days in vitro* in a raw. Such monitoring time windows are almost 10 times longer than what is possible with commercial techniques and more than double than what shown on alternative cutting-edge technologies (Jahed et al. 2022).

To demonstrate the performances for cardiotoxicity screening, we also measure how a known compound, pentamidine, affects cardiac ion channels over long-term exposure. As expected, the molecule does not affect the AP waveforms after acute administration. However, we can correctly observe the effects on the cardiac cells as exposure time increases up to hundreds of hours and the cardiomyocytes recovery during the subsequent drug washout period. Furthermore, given the importance of chronic cardiotoxicity in the development of new anticancer drugs, we further tested our approach by monitoring the effects of repeated administration of doxorubicin on the electrophysiological activity of hiPSC-CMs for an extremely long period. Finally, we demonstrate that the long-term assessment of APs by consecutive optoporation sessions has no effects on cell health.

Our results demonstrate that optoporation may be a valuable tool to develop reliable cardiac cellular models for therapeutic and diagnostic applications and to assess the delayed drug-induced cardiotoxicity during preclinical phases of drug development process.

## Materials and Methods

### hiPSC cardiomyocytes culture on commercial MEAs

HiPSC-derived cardiomyocytes were purchased from Cellular Dynamics International. iCell cardiomyocytes, a mix of ventricular, atrial and nodal-like cells, were directly plating on 60-6wellMEA200/30iR-Ti-rcr (Multichannel Systems) at a density of 16,000 cells per well and grown according to the specifications of the commercial supplier. MCS MEAs were coated with fibronectin for 1h at 37°C, 5% CO_2_ in a humidified incubator to promote the cells adhesion. Electrophysiological recordings were performed starting from 9 days post-plating and repeated every 2-3 days.

### Chemical compounds

Pentamidine and doxorubicin were purchased from Sigma-Aldrich. The 10 mM stock solutions were prepared in dimethyl sulfoxide (DMSO). Drug dilutions were performed in pre-warmed (37 °C) iCell-Maintenance Medium.

### Electrophysiology recording

All recordings from MCS-MEAs were performed at 37 °C outside the incubator. The cell medium from each sample was changed two hours before the measurements. All data sets were analyzed by Matlab software. MEA recordings were obtained with a MEA2100-Mini-System from the company Multi Channel Systems MCS GmbH. Recordings with the MEA2100-Mini-System were acquired at 20 kHz acquisition frequency, high-pass hardware filtering of 0.1 Hz, and low-pass hardware filtering of 10000 Hz.

### Immunofluorescence staining

After six consecutive intracellular recording sessions (21 DIVs), for immunofluorescence staining, the cells were fixed with 4% (w/v) formaldehyde in phosphate-buffered saline (PBS), permeabilized with 1% saponin (15 min), blocked with 3% bovine serum albumin (BSA) for 30 min followed by incubation with TNNT2 (mouse anti-TNNT2 A25969) antibody diluted 1:1000 in a solution of 3% BSA for 3h at room temperature. After three washes in PBS, cells were incubated with Alexa Fluor 488 donkey anti-mouse secondary antibody (diluted 1:250 in blocking solution) for 1h at room temperature, followed by three washes with PBS. Finally, cell nuclei were counterstained with DAPI cells and then the images were acquired using an inverted Nikon microscope by using filters for Alexa 488 channel (Ex/Em 495/519 nm) and DAPI channel (Ex/Em 358/461 nm).

### Laser optoporation

Laser pulse trains were applied on the surface of MCS MEAs to porate hiPSC-CMs. For laser poration, the first harmonic (*λ* = 1064 nm) of a Nd:YAG (neodymium:yttrium–aluminium–garnet) solid-state laser (Plecter Duo (Coherent)) with an 8 ps pulse width and 80 MHz repetition rate was used as radiation source, with an average power of approximately 9 mW after the objective. The laser was coupled to an inverted optical setup able to accommodate the acquisition system MEA2100-Mini from Multi Channel Systems MCS Gmbh. A 50X air objective (NA = 0.45, working distance = 25 mm) was used during the experiments to observe the cells on the devices and to focus the NIR laser used for poration.

### Data and statistical analysis

A custom-made algorithm was specifically developed in MATLAB (The Mathworks, Natick, MA, USA) to perform data analysis. The algorithm allows for accurate characterization of the intracellular action potentials waveforms, the intracellular APs duration at different amplitude values with respect to the maximum amplitudes (30%, 50% and 90%), and the beating rate. Data were expressed as mean ± standard error of the mean (SEM). Statistical analysis was performed to determine significant differences between all sample pairs by using MATLAB. Since data do not follow a normal distribution (evaluated by the Kolmogorov-Smirnov normality test), a non-parametric Mann-Whitney U-test was used. In all cases, significance was defined as p ≤ 0.05.

## Results

### Long-term monitoring of APs from hiPSC-CMs

We cultured hiPSC-CMs on 60-electrode 6-well commercial MEAs (Fig.1a, b). The employed commercial MEA presents transparent titanium nitride (TiN) electrodes that allow for applying the laser-based optoporation protocol from the bottom of the device (Fig. 1c) by means of an inverted optical setup. Thanks to the electrode transparency, laser radiation from the bottom transmits through the electrode and reaches the electrode-cell interface, where it generates charge ejection from the electrode material and leads to localized poration of the cellular membrane (Melle et al. 2020). Optoporation from the backside of the MEA devices is fundamental for preserving a perfect sealing of the device during recordings, helping maintaining the necessary sterility for long-term measurements. We performed optoporation and intracellular AP recording starting after approximately 9 *days in vitro* (DIVs) and we repeated the measurements every 2-3 days up to a maximum of 44 DIVs, for a total of more than 30 monitoring days (Fig. 1d). After this time window, cardiac cells started to show detachment from the MEA surface and measurements became unreliable. In each measurement day, we recorded intracellular APs from all 9 electrodes in each well of the MEA obtaining the mean AP waveform of each cardiac monolayer during a long period of time. Fig.1e depicts exemplary mean APs at different DIVs from the same cardiac monolayer (the same well in the 6-well MEA). The AP waveforms present high signal-to-noise ratio and allow us for evaluating transmembrane ion currents from the same syncytium over extremely long periods. Furthermore, in order to evaluate the effect of repeated optoporation on cardiomyocytes health, immunofluorescence for cardiac troponin-T (cTnT) was performed on hiPSC-CMs after six consecutive intracellular recording sessions. Immunofluorescence images highlights the typical organization and filament-like structure of Troponin T of cardiomyocytes after repeated optoporation, demonstrating that the long-term assessment of APs has no significant effects on the cell health (Fig. 1f). Thus, our approach permits the long-time assessment of APs preserving the health of the cardiac cells.

**Fig. 1:**
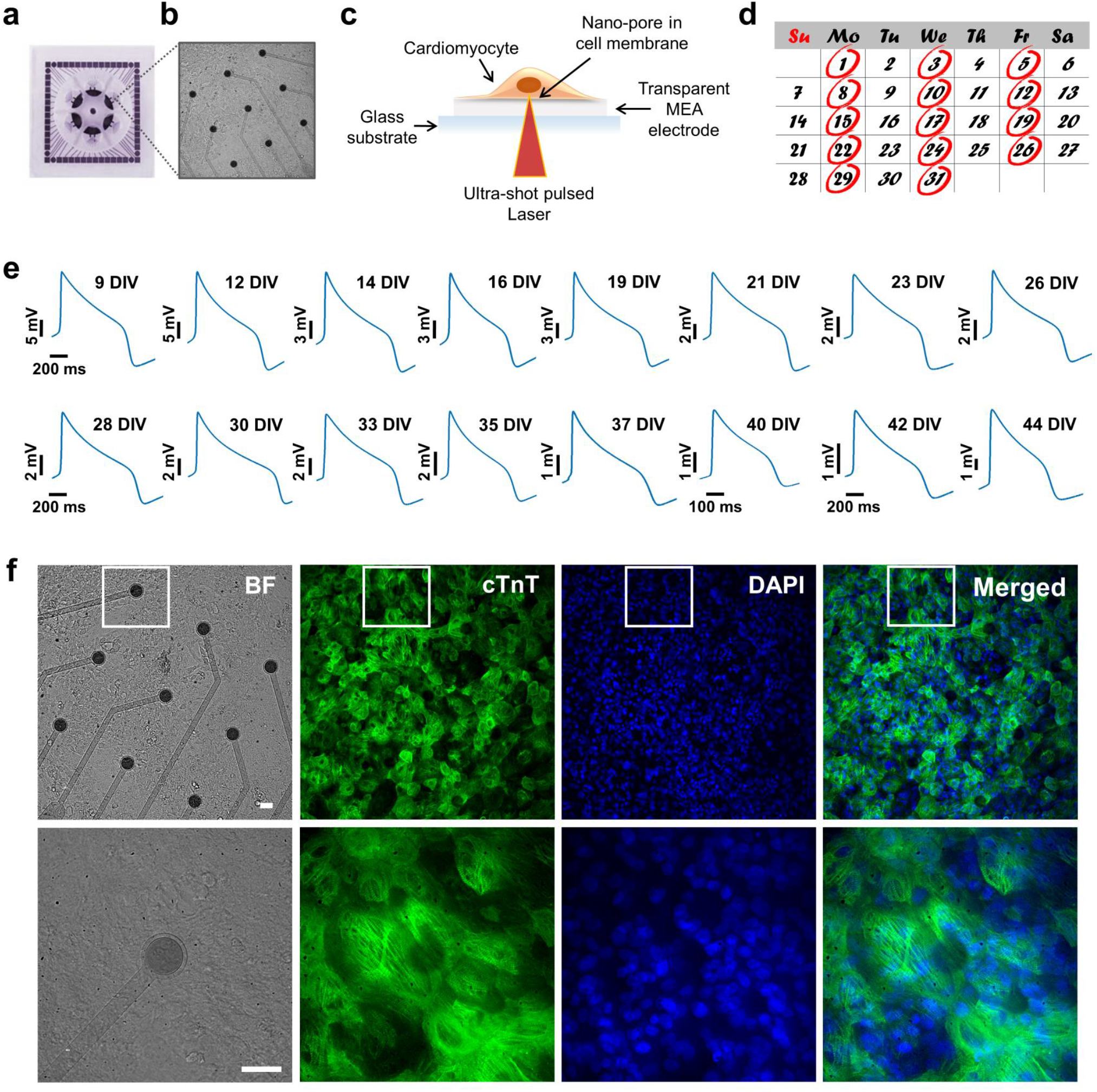
Long-term intracellular AP recordings from hiPSC-CMs. (**a**) Photograph of a 6-well MEA (60-6wellMEA200/30iR-Ti-rcr). (**b**) hiPSC-CMs at 9DIVs in a single MEA well. (**c**) Representative scheme of laser-based optoporation on 6-well MEA devices. (**d**) Time course of APs recording. (**e**) Action potential mean waveforms of cardiac syncytium in a well of 6-well MEA at different DIVs. (**f**) Immunofluorescence images at 20X magnification of hiPSC-CMs on 6-well MEA after 6 repeated opotoporation procedures and at 60X magnification of the area highlighted with white box. Scale bar: 30 µm.

Furthermore, our analysis of AP duration at different amplitudes (30% (APD30), 50% (APD50) and 90% (APD90)) of repolarization shows variations of APD and beating rate over days, remaining though within the physiological range of 600-800 ms (Bett et al. 2013; Sala et al. 2017; Mannhardt et al. 2020) (Fig. 2a, b). These results indicate that hiPSC-CMs continue to mature during time in culture. Hence, long-term assessments of cardiotoxicity should consider this behavior when evaluating drug effects on ion channels.

**Fig. 2:**
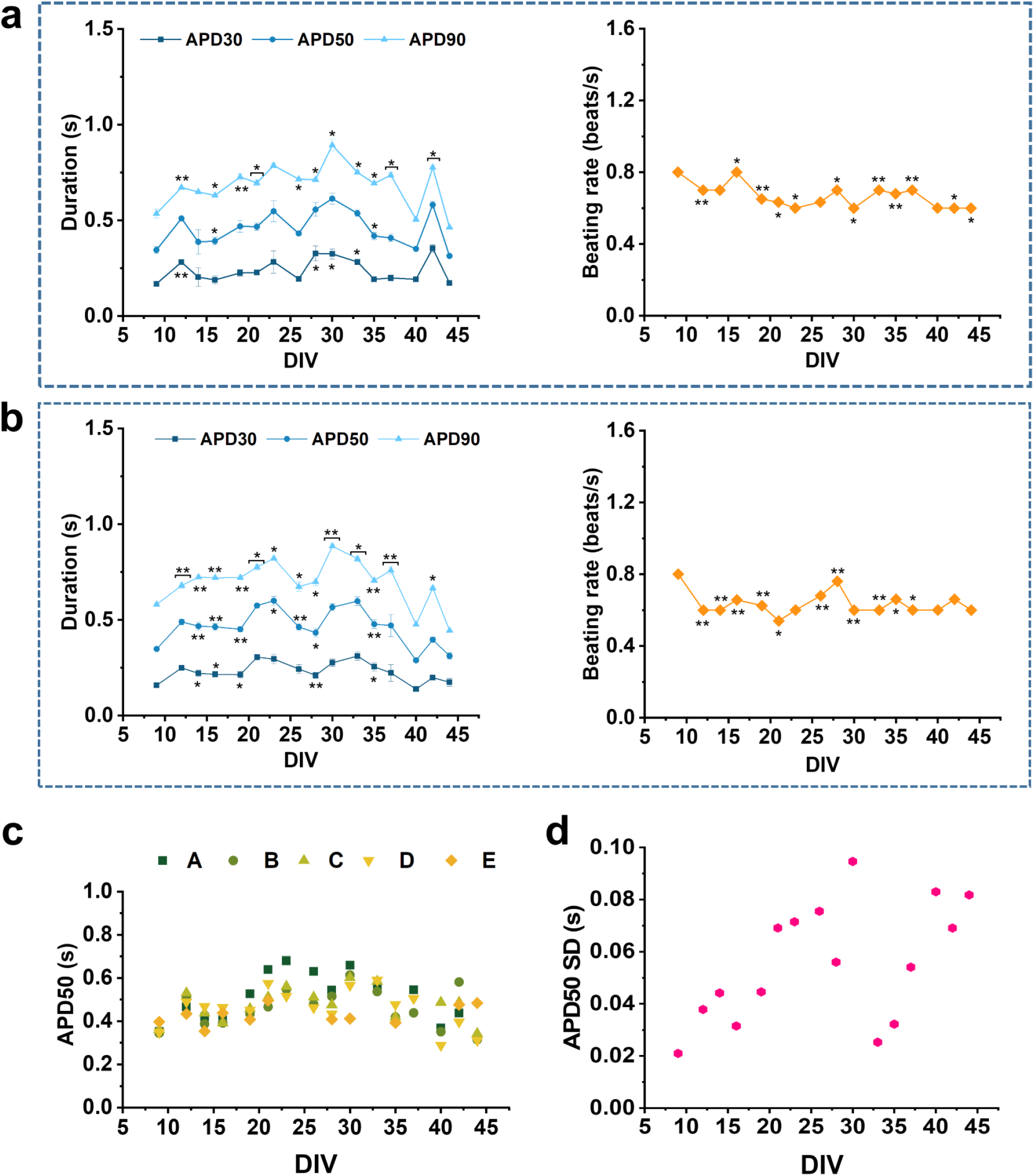
Long-term intracellular AP recordings from hiPSC-CMs. (**a, b**) Action potential duration at different amplitudes (APD30, 50, 90) and beating rate for two different wells of a 6-well MEA. (**c**) Comparison of the APD at 50% repolarization (APD50) of each well over time. (**d**) Standard deviation of APD50 among wells. Data are represented as mean ± SEM. P-values are calculated using Mann-Whitney U-test, *0.01<P<0.05, **0.001 <P<0.01, ***0.0001 <P<0.001, ****P<0.0001.

In figure 2c, we report APD50 values of 5 different wells in a 6-well MEA of the same cell culture preparation from 9 to 44 DIVs. At 9 DIVs, we observe that APD50 values are relatively similar as it is expected because cells have been treated exactly in the same way and had a short time to diverge. As the cells remain longer in culture, we observe that APD50 values start to diverge among wells. This is highlighted in figure 2d where we present the standard deviation (SD) of APD50 among the wells along the whole culture time. It can be observed that SD tends to increase over time. We suggest this to be an important factor to consider when assessing long-term cardiotoxicity, because the comparison between independent syncytia could lead to wrong conclusions about chronic drug effects. More reliable results can be indeed obtained by testing drugs within the same syncytium, without using a different culture well as control.

### Long-term AP monitoring from single cells

Our approach for long-term AP recordings not only represents a valuable tool for long-term assessment of large hiPSC-CM syncytia, but it also enables long-time monitoring of APD from single cardiac cells in each well of 60-electrode 6-well commercial MEAs (Fig. 3a). Fig. 3b shows AP at different DIVs from the same cardiac cell on one electrode. The AP duration analysis at different amplitudes (APD30, APD50 and APD90) of repolarization from two single electrodes in two different wells highlights the variations of APD over days (Fig. 3c, d). Such feature results from the extremely low invasiveness of cell optoporation, which allows for porating and measuring the same cell for several times without negative effects.

**Fig. 3:**
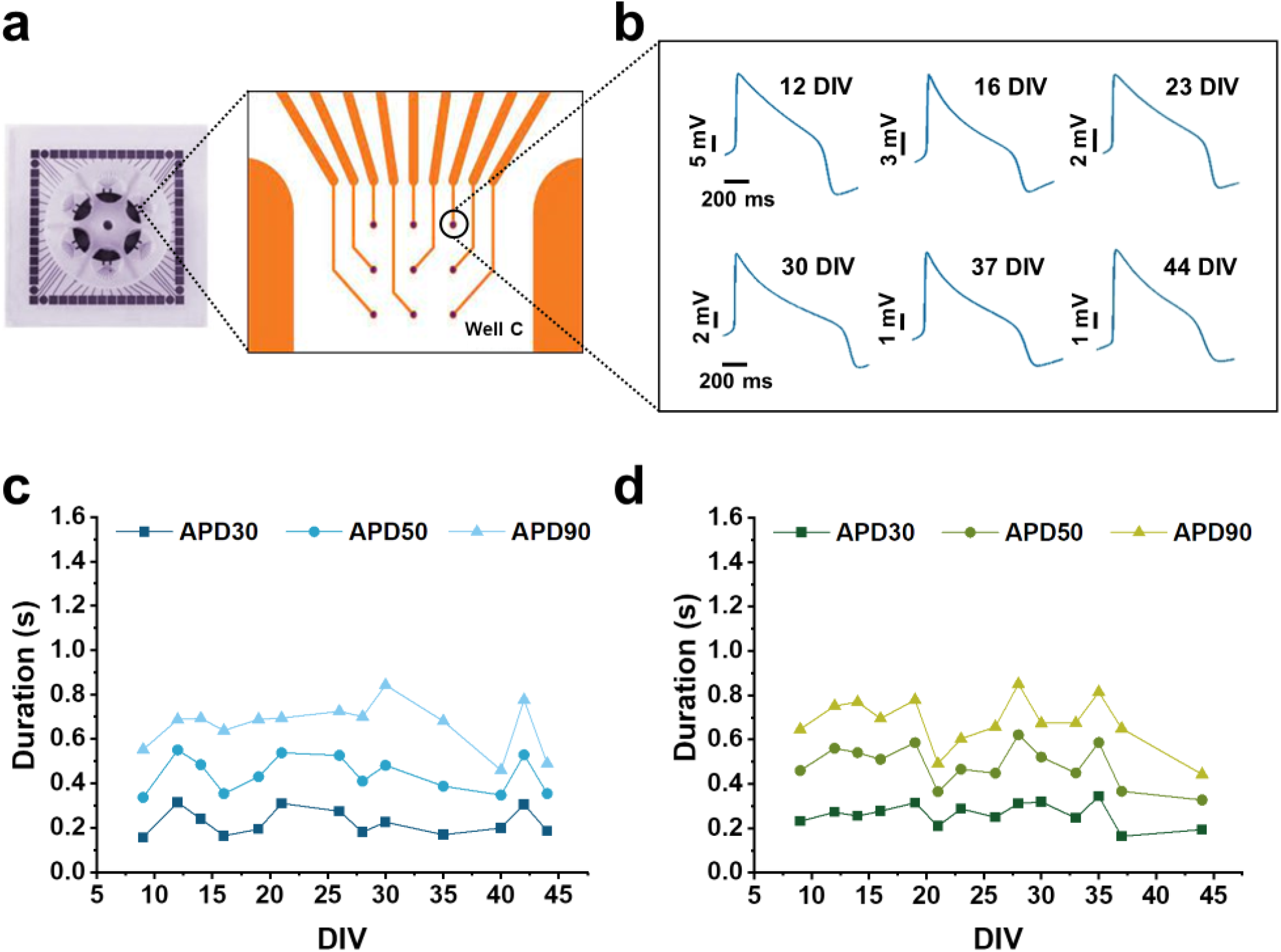
Long-term intracellular AP recordings from single cardiac cells. (**a**) Photograph of a well of 6-well MEA (60-6wellMEA200/30iR-Ti-rcr). (**b**) Action potential waveform of single cells in a well of a 6-well MEA at different DIVs. (**c, d**) Action potential duration at different amplitudes (APD30, 50, 90) from two single electrodes in two different wells.

The monitoring of changes in cardiac transmembrane ion currents of single cells in large syncytia is fundamental for evaluating cell-specific drug effects. Indeed, inhomogeneous drug distribution (Pinto and Howell 2007) and/or phenotype differences can influence the electrical response of cells to the same drug.

Furthermore, despite the progresses in development of subtype-specific differentiation protocols (Zhang et al. 2011; Devalla et al. 2015; Birket et al. 2015), obtaining pure populations of ventricular-, atrial-, or nodal-like hiPSC-CMs is still challenging. Following the AP evolution of single cardiac cells may help the electrophysiological classification of hiPSC-CMs into cardiac subtypes based on their AP profiles, as this is currently the gold standard for cardiomyocytes characterization (Zhang et al. 2009; Lieu et al. 2013). However, the existing approaches to assess the AP profiles make the cells unviable for subsequent analysis preventing the proper monitoring of maturation of the cardiomyocytes over time. Our approach may overcome this limitation enabling the monitoring in time of hiPSC-CM maturation, facilitating the development of more reliable and predictive cardiac cellular models intended for therapeutic and diagnostic applications.

### Chronic drug effects on hERG channel

In order to assess the ability of our approach to detect long-term cardiotoxic drug effects, we tested a hERG trafficking inhibitor, pentamidine, which produces adverse cardiac effect after several hours of continuous exposure (Asahi et al. 2019). Pentamidine is used to treat several parasitic diseases but it is also associated with drug-induced long QT syndrome and Torsades de Pointes (TdP) (Obergrussberger et al. 2016). Interestingly, this drug has no direct hERG-blocking activity but inhibits the hERG trafficking to the cell membrane, reducing its expression on the cell surface (Cordes et al. 2005; Nogawa and Kawai 2014). This mechanism of action may go undetected in most conventional cardiac safety assays (i.e. conventional or automated patch clamp), highlighting the necessity for longer-term assays to evaluate indirect effects on the ion channels.

We monitored APs of the same hiPSC syncytia up to 11DIVs (from 14 DIVs to 23 and 25 DIVs), exposing cells to pentamidine for 3-4 DIVs and then monitoring cell recovery after drug washout. We tested pentamidine at 0.5 µM and 1.5µM concentrations. Fig. 4a, b depict AP waveforms along the whole 11 DIVs experiment at the two different concentrations. In fig. 4c and 4d, we summarize the effects by tracing APD30, APD50 and AP90 over time. Pentamidine shows both concentration- and time-dependent effects after long exposure time. At both tested concentration, pentamidine leads to APD increase starting from 24 hours of incubation and is strengthened in the following incubation days (Fig. 4c, d). At higher concentration of pentamidine, the AP elongation is greater and occurs early compared to the lower concentration and the effect can still be observed 1 day after drug washout (Fig. 4d). Furthermore, we checked whether the cardiac monolayer had the ability to spontaneously recover from pentamidine effect by monitoring the APD after the drug washout. Our results show a return to the APD physiological range for both tested concentration, this occurs the day after drug washout for the lower concentration and two days after drug washout for the higher concentration of pentamidine administrated (Fig. 4c, d). Finally, we observe also beating rate reduction induced by pentamidine in a concentration-dependent manner (Fig. 4e, f).

**Fig. 4:**
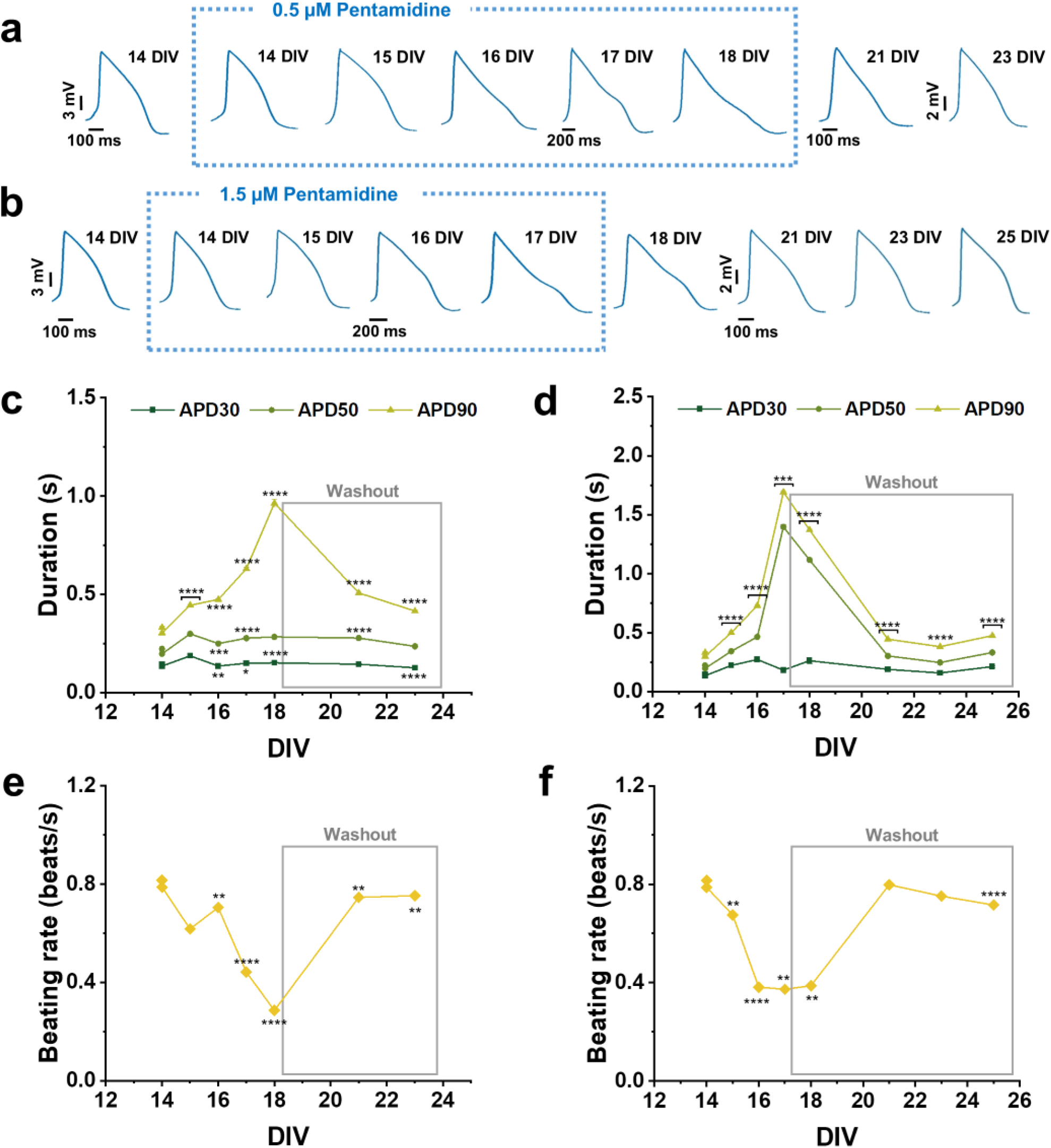
Long-term effect of pentamidine on hiPSC-CMs. (**a, b**) Action potential mean waveforms at different days after 0.5 µM and 1.5 µM pentamidine administration, respectively. (**c, d**) Action potential duration at different amplitudes (APD30, 50, 90) after 0.5 µM and 1.5 µM pentamidine administration, respectively. (**e, f**) Beating rate after 0.5 µM and 1.5 µM pentamidine administration, respectively. Data are represented as mean ± SEM. P-values are calculated using Mann-Whitney U test, *0.01 < P < 0.05, **0.001 < P < 0.01, ***0.0001 < P < 0.001, ****P < 0.0001.

### Long-term effects of doxorubicin on cardiomyocytes

We have also explored the advantages of our approach in detecting long-term effects of doxorubicin on cardiac ion channels. Doxorubicin is an anthracycline antibiotic widely used in the treatment of several cancers (Todaro et al. 2013). However, doxorubicin causes chronic cardiotoxicity that can arise as late as 10 years after last administration, making extremely challenging to predict it during acute preclinical studies (Franco and Lipshultz 2015; Dong and Chen 2018). Therefore, to predict the long-term cardiotoxicity of this kind of drug, it is necessary to increase the assessment time window. Currently, the long-term doxorubicin cardiotoxicity has been explored by endpoint assays using xCELLigence Real Time Cell Analyser (ACEA Biosciences) to evaluate cytotoxicity and beating frequency of hiPSC-CMs and the transcriptomic analysis to identify genomic biomarkers for anthracycline-induced cardiotoxicity (Chaudhari et al. 2016). Furthermore, chronic doxorubicin-induced cardiotoxicity was also assessed using the standard cell viability assay (MTT 3-(4,5-dimethylthiazol-2-yl)-2,5-diphenyltetrazolium bromide) (Karhu et al. 2020), investigating the metabolite signatures (Chaudhari et al. 2017), the contractile motion properties and cardiac biomarker development (Kopljar et al. 2017). However, all current approaches for assessment of chronic doxorubicin-induced cardiotoxicity are based on endpoints assays and do not allow for a continuous monitoring of transmembrane ion currents to evaluate their role in delayed cardiac adverse effects. On the contrary, optoporation on transparent MEAs enables the monitoring of APs of the same hiPSC syncytia up to 24DIVs (from 9 DIVs to 33 DIVs) after repeated doxorubicin exposure and the evaluation of cell recovery after drug washout. Fig. 5a, c depict APs waveforms along the entire 24 DIVs experiment at the two different concentrations (5 nM and 10 nM). We observed a decrease in spike amplitude for both tested concentrations (Fig. 5a, c), an increase in beating rate for lowest tested concentration of doxorubicin and a decrease for the highest tested concentration only at some DIVs (Fig. 5b, d). We detected that the prolonged doxorubicin treatment induces a decrease of APD90 at several DIVs for both tested concentrations as observed in the common trend visible in all wells (Fig. 5e, 5f). Moreover, we observed no spontaneous recovery from doxorubicin effects after drug washout. Our electrophysiological data are comparable with Maillet et al. studies, in which they observed both acute and chronic effects of doxorubicin in hiPSC-CMs, including decrease in spike amplitude, increase in beat rate and shortening in corrected field potential duration (cFPD) after 20 hours of treatment but using concentrations of doxorubicin in micromolar range (Maillet et al. 2016). However, despite the electrical activity of hiPSC-CMs is noticeably affected only by concentrations of doxorubicin in the micromolar range, nanomolar concentrations are already enough to affect cell viability and cause mitochondrial alterations (Louisse et al. 2017). Moreover, arrhythmic beating activity and decreased beating amplitude were also reported upon a 6 day exposure to doxorubicin (150 nM), possibly correlated to a doxorubicin-induced decrease in cell number or structural cardiomyocyte changes (Chaudhari et al. 2016). In this context, our results extend the time window of assessment of long-term cardiotoxicity cause by doxorubicin by evaluating an accurate functional parameter such as the action potential, thus improving the understanding of the complex mechanisms underlying anthracycline cardiotoxicity.

**Fig. 5:**
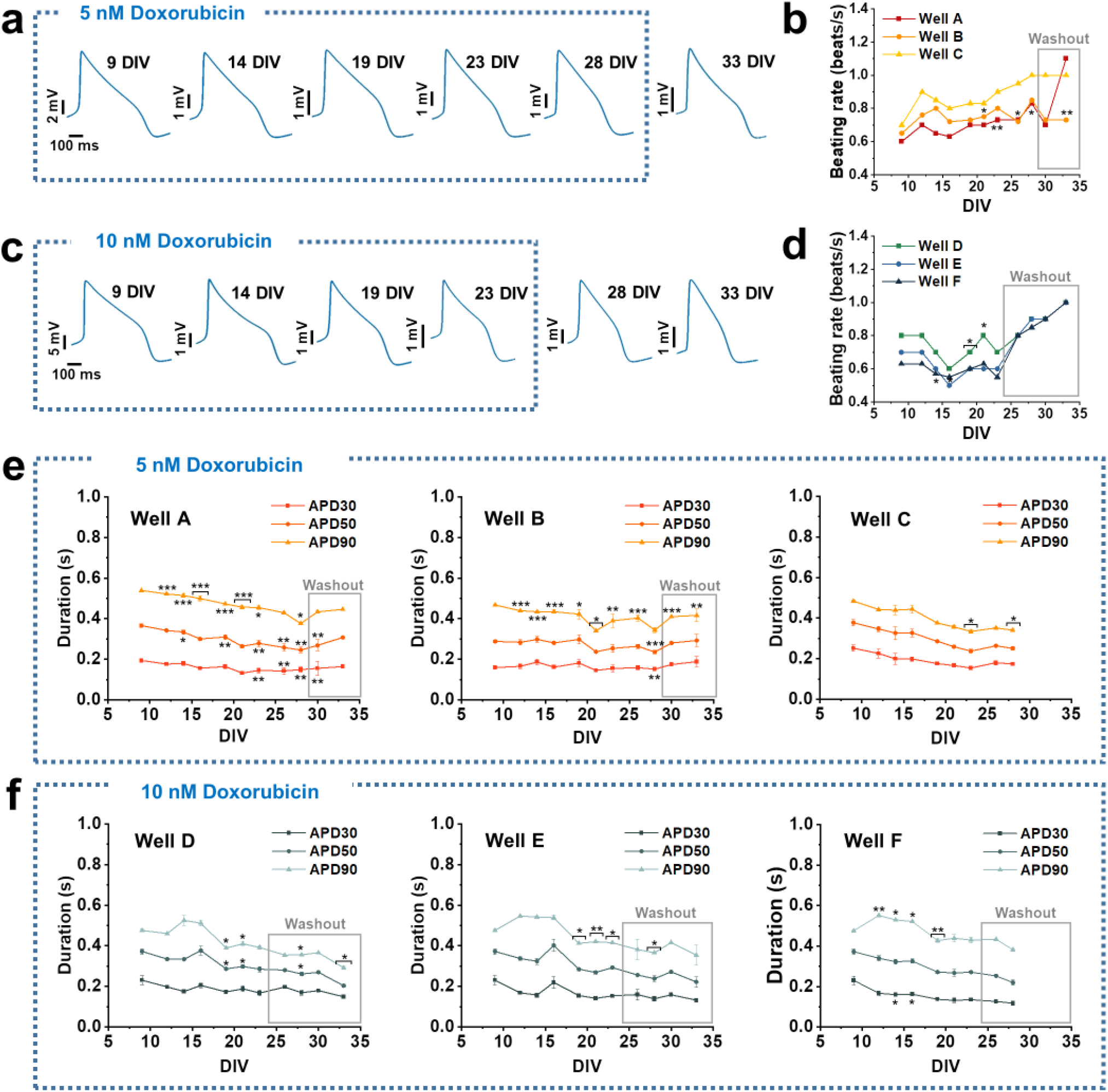
Long-term effect of doxorubicin on hiPSC-CMs. (**a, b**) Action potential mean waveforms at different days after 5 nM and 10 nM doxorubicin administration, respectively. (**b, d**) Beating rate after 5 nM and 10 nM doxorubicin administration in different wells. (**e, f**) Action potential duration at different amplitudes (APD30, 50, 90) after 5 nM and 10 nM doxorubicin administration, respectively. Data are represented as mean ± SEM. P-values are calculated using Mann-Whitney U test, *0.01 < P < 0.05, **0.001 < P < 0.01, ***0.0001 < P < 0.001, ****P < 0.0001.

## Discussion

In this work, we detect the effects of compounds to the APs of hiPSC-CMs and we demonstrate long-term cardiotoxicity assessment capabilities by performing optoporation and intracellular AP recordings for up to 35 DIVs on the same cardiac cells. We obtained data on multiwell MEAs with transparent electrodes from large hiPSC-CM syncytia that represent excellent biological models for cardiotoxicity assessments. The investigated duration on the same biological samples is substantially longer than what standard screening technologies offer. Hence, the approach promises to improve drastically the detection of chronic toxic effects of drug candidates.

Our analyses on APD changes over time among different culture wells highlight the risks of using independent syncytia as reference for dose-response measurements, because cardiac cell cultures can present substantial differences during further maturation in long-term experiments. Hence, the possibility to follow AP changes over long-time on the same cells or same syncytium represent an efficient way to increase the reliability of the tests.

Our approach not only enables the accurate electrophysiological assessment of hiPSC-CMs maturation over long time, but also the monitoring of changes in cardiac transmembrane ion currents of single cells in large syncytia, representing an advantage for evaluating cell-specific drug effects and/or phenotype characterization of cardiac culture. Considering the impact of the cardiac model choice and the maturation state of hiPSC-CMs on drug compound responsiveness (da Rocha et al. 2017; Karhu et al. 2020), optoporation may significantly contribute to establish an appropriate experimental model for delayed toxicity screening in early drug development.

In details, we correctly detected for 11 DIVs (from 14 DIVs to 25 DIVs) alterations of cardiac transmembrane ion currents due to the chronic administration of pentamidine, an hERG trafficking inhibitor which only induces cardiotoxic effects after longer term exposure (Kuryshev et al. 2005; Obergrussberger et al. 2016; Asahi et al. 2019). We observed the prolongation of APD after long-term exposition to pentamidine and the ability of cardiomyocytes to recover spontaneously after drug washout. Furthermore, since cardiotoxicity is one of the major issues of anti-cancer therapy, we also investigated the capabilities of optoporation to assess the effects of long-term administration of doxorubicin, the most widely used chemotherapeutic drug for cancer therapy (Sritharan and Sivalingam 2021). We monitored APs of the same hiPSC syncytia up to 24 DIVs (from 9 DIVs to 33 DIVs) after repeated doxorubicin exposure finding a decrease in spike amplitude for both tested concentrations and an increase in beating rate only for the lower concentration. We found a decrease in APD90 at several DIVs for both tested concentrations. Moreover, we observed that cardiomyocytes were not able to spontaneously recover from doxorubicin after washout.

Taken together, our results demonstrate that optoporation may facilitate the development of more reliable cardiac cellular models intended for therapeutic and diagnostic applications thanks to the ability to monitor hiPSC-CMs maturation in terms of transmembrane ion currents. Furthermore, our method may be an effective *in vitro* strategy to assess the long-term exposure of hiPSC-CMs to compounds and delayed drug-induced cardiotoxicity in general, drastically reducing the attrition rates and market withdrawals during the drug development process.

## Author Contributions

Giuseppina Iachetta: Methodology, Formal analysis, Investigation, Data curation, Writing – original draft, Writing – review & editing. Giovanni Melle: Methodology, Data curation. Nicolò Colistra: Methodology, Data curation. Francesco Tantussi: Methodology. Francesco De Angelis: Conceptualization, Writing – review & editing, Supervision. Michele Dipalo: Conceptualization, Methodology, Supervision, Investigation, Writing – original draft, Writing – review & editing.

